# Exploring the role of glycans in the interaction of SARS-CoV-2 RBD and human receptor ACE2

**DOI:** 10.1101/2021.03.30.437783

**Authors:** Kien Nguyen, Srirupa Chakraborty, Rachael A. Mansbach, Bette Korber, S. Gnanakaran

## Abstract

COVID-19 is a highly infectious respiratory disease caused by the novel coronavirus SARS-CoV-2. It has become a global pandemic and its frequent mutations may pose new challenges for vaccine design. During viral infection, the Spike RBD of SARS-CoV-2 binds the human host cell receptor ACE2, enabling the virus to enter the host cell. Both the Spike and ACE2 are densely glycosylated, and it is unclear how distinctive glycan types may modulate the interaction of RBD and ACE2. Detailed understanding of these determinants is key for the development of novel therapeutic strategies. To this end, we perform extensive all-atom simulations of the (i) RBD-ACE2 complex without glycans, (ii) RBD-ACE2 with oligomannose MAN9 glycans in ACE2, and (iii) RBD-ACE2 with complex FA2 glycans in ACE2. These simulations identify the key residues at the RBD-ACE2 interface that form contacts with higher probabilities, thus providing a quantitative evaluation that complements recent structural studies. Notably, we find that this RBD-ACE2 contact signature is not altered by the presence of different glycoforms, suggesting that RBD-ACE2 interaction is robust. Applying our simulated results, we illustrate how the recently prevalent N501Y mutation may alter specific interactions with host ACE2 that facilitate the virus-host binding. Furthermore, our simulations reveal how the glycan on Asn90 of ACE2 can play a distinct role in the binding and unbinding of RBD. Finally, an energetics analysis shows that MAN9 glycans on ACE2 decrease RBD-ACE2 affinity, while FA2 glycans lead to enhanced binding of the complex. Together, our results provide a more comprehensive picture of the detailed interplay between virus and human receptor, which is much needed for the discovery of effective treatments that aim at modulating the physical-chemical properties of this virus.

## INTRODUCTION

The severe acute respiratory syndrome coronavirus 2 (SARS-CoV-2) is responsible for the highly contagious coronavirus disease 2019 (COVID-19). It has led to an ongoing global pandemic where emerging mutations may require continuous development of novel therapeutic strategies. Similar to other coronaviruses, SARS-CoV-2 uses the Spike glycoprotein to interact with human host cells during viral infection [1,2]. Specifically, the receptor binding domain (RBD) of the Spike glycoprotein binds the angiotensin-converting enzyme 2 (ACE2) receptor on the host cell (Figure 1). This facilitates fusion of virus and host cell membranes, leading to entry of the virus into the cell.

**FIG. 1.**
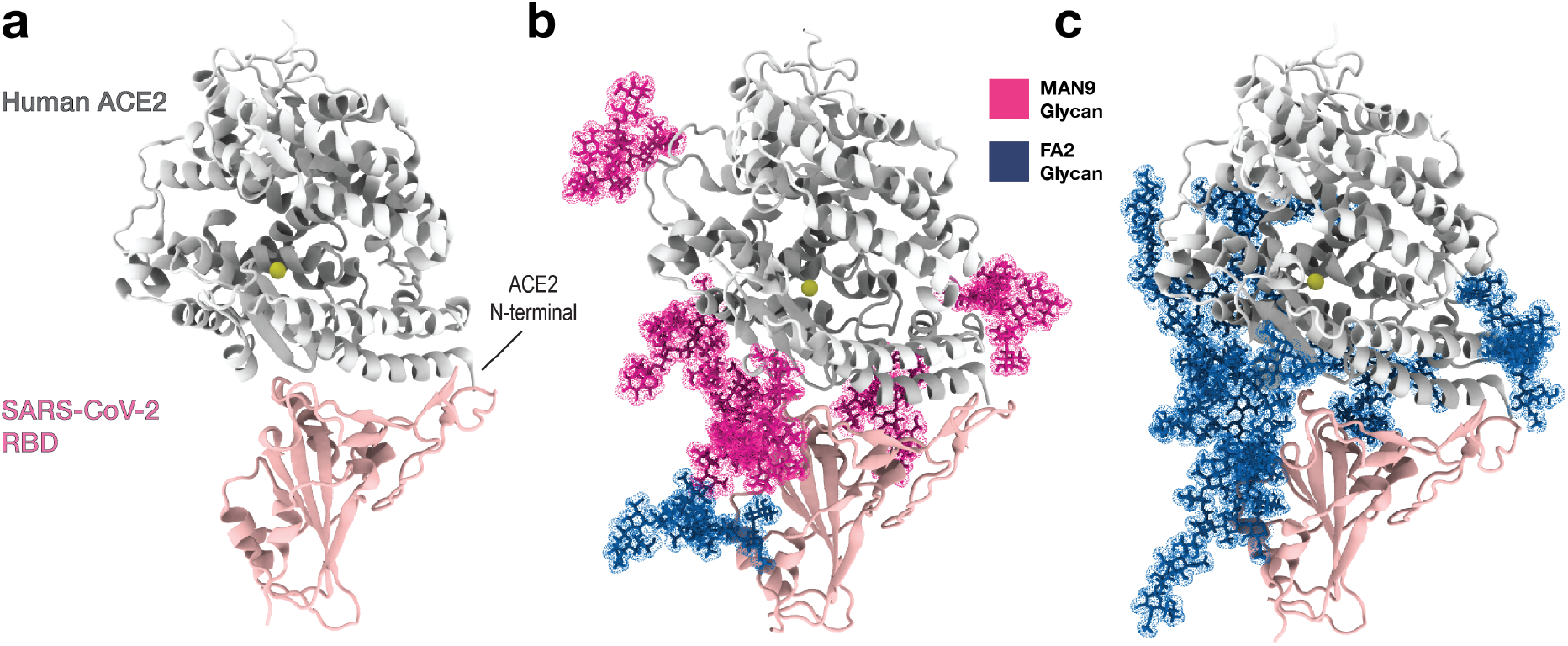
Simulations of the RBD-ACE2 complex in the absence and presence of different glycan types. **(a)** Structural representation of human receptor ACE2 (silver) bound to RBD of SARS-CoV-2 (pink). The yellow bead in ACE2 depicts the zinc ion that is coordinated by His374, His378, Glu402, and one water molecule. **(b)** The protein complex of panel (a) is shown with six MAN9 glycans (magenta) bound to ACE2 (at Asn53, Asn90, Asn103, Asn322, Asn432, Asn546) and one FA2 glycan (blue) bound to RBD (at Asn343). **(c)** Same complex as in (b) except that the six ACE2 glycans are FA2. Panels (a), (b), and (c) represent three systems for which separate sets of simulations were performed. All molecular graphics were created using VMD [47]

Since RBD binding to ACE2 is key to viral entry, this interaction is a major target for development of antibody therapeutics and vaccine design. Mutations in the RBD can alter virus-receptor interaction [3, 4], and thus, viral infectivity. Recent structural studies have revealed the residue-residue contacts between SARS-CoV-2 RBD and ACE2, suggesting possible interactions that may determine the stability of the complex [5–7]. However, from these static data, it is unclear which of these residues are the critical ones that form interactions most frequently. While a recent computational study has addressed this question [8], their results are based on relatively short simulations (~ 5 μs) that may not achieve accurate statistics. In addition, both the viral Spike and host receptor ACE2 are densely glycosylated with asparagine linked N-glycans [9, 10]. Since some of these glycans are spatially located near the RBD-ACE2 interface (Figure 1), it is necessary to assess how they may affect the binding affinity of RBD to ACE2. Previous experiments have elucidated some roles of glycans where they, for instance, can impact antibody interactions and epitope exposure [11]. Furthermore, different glycan types with characteristic structures can be critical for pathogen-host interaction [12, 13]. Other experiments have shown that disruption of ACE2 glycosylation can impair SARS-CoV-1 viral entry into cells [14]. However, due to structural complexity and heterogeneity of glycans, together with limited instrumental sensitivity [15], it is unclear how individual glycan types could distinctly modulate RBD binding. Such thorough understanding is needed to suggest more precise strategies for future experiments that seek to design effective treatments. For this purpose, we apply extensive molecular dynamics (MD) simulations (> 200 μs) that elucidate the detailed interplay between SARS-CoV-2 RBD and host ACE2, and the role of different glycan types during this process. Specifically, we study RBD-ACE2 complexes that either have a) mannose-9 (MAN9) glycans attached to six ACE2 residues: i.e. Asn53, Asn90, Asn103, Asn322, Asn432, and Asn546 or b) sialated complex flucosylated 2-antennae (FA2) glycans attached to the same afore-mentioned ACE2 residues. In both of these glycan-included models, the Spike RBD is built with a single FA2 glycan at Asn343 (see SI Methods for details).

To dissect the determinants of RBD-ACE2 binding, we performed all-atom explicit-solvent simulations of three systems: (i) the RBD-ACE2 complex without glycans (Figure 1a), (ii) RBD-ACE2 with six MAN9 glycans on ACE2 and one FA2 glycan on RBD (Figure 1b, S1), and (iii) RBD-ACE2 with six FA2 glycans on ACE2 and one FA2 glycan on RBD (Figure 1c). These three separate sets of simulations comprise of 90 trajectories that have a combined simulated time of more than 225 μs. Statistical analysis of these long simulations reveal the key residues of RBD and ACE2, which form binding contacts with higher probabilities. This provides a complementary view to structural data by quantifying the significance of identified interactions. Importantly, we find that this RBD-ACE2 contact signature is not affected by the presence of MAN9 or FA2 glycans, suggesting RBD-ACE2 contacts are inherently robust. Using our simulated interactions as foundation, we evaluate the effect of N501Y, a recent mutation that accounts for the majority of new infections in South East England. Specifically, we show that N501Y in RBD introduces additional stabilizing interactions with Y41 and K353 of ACE2, which can contribute to enhanced infectivity of the new strain. Furthermore, we show that the glycan on Asn90 of ACE2 (regardless whether MAN9 or FA2) contacts the RBD with distinctly higher probabilities, compared to the remaining glycans on ACE2. This indicates that the Asn90 glycan may play a critical role in binding/unbinding events of RBD, as previously suggested by atomic force microscopy (AFM) experiments [16], biolayer interferometry [17], and other biochemical assays [18]. Finally, an analysis of binding energetics demon-strates that each glycan type is associated with distinct enthalpic contributions to binding affinity. Specifically, we show that MAN9 glycans on ACE2 lead to weaker RBD-ACE2 binding, while the FA2 glycans, with their negatively charged sialic acid tips, stabilize the complex. Together, the results provide a quantitative framework that allows future studies to more precisely modulate the infectivity of SARS-CoV-2 and improve immunogen design.

## RESULTS

### Contacts between ACE2 and RBD are not altered by the presence of glycans

To obtain a detailed picture of how SARS-CoV-2 RBD interacts with human receptor ACE2, it is necessary to identify the residues that form close contacts at the RBD-ACE2 interface. Furthermore, we investigate how this RBD-ACE2 interaction is affected by different glycan species that are present on the ACE2 surface. To this end, we performed three separate sets of all-atom explicit-solvent simulations of (i) the RBD-ACE2 complex without glycans (Figure 1a), (ii) RBD-ACE2 with MAN9 glycans on ACE2 (Figure 1b), and (iii) RBD-ACE2 with FA2 glycans on ACE2 (Figure 1c). For each simulation set, we calculated the probability of ACE2 and RBD residues forming contacts with each other (Figure 2a-c). Here, a contact is considered formed if the smallest distance between the heavy atoms of two amino acid residues is within 4 Å.

**FIG. 2.**
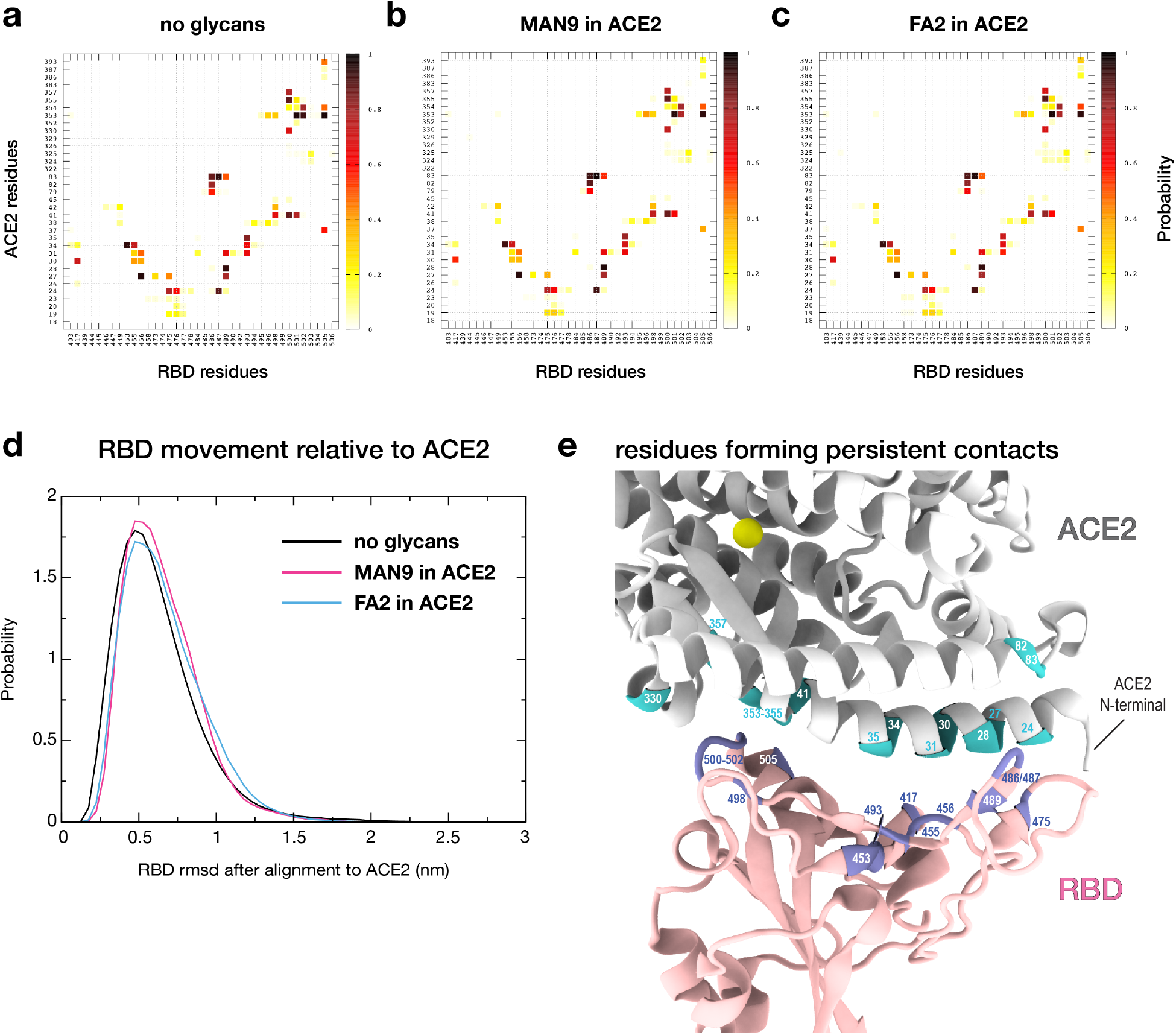
Contact signature of ACE2 and RBD is robust to the presence and types of glycans. **(a)** Probability of contact formation between ACE2 and RBD residues. This contact map was calculated from the simulations where glycans are not present (cf. Figure 1a; see Table S1 for list of contacts). To elucidate the role of glycans, we repeated this analysis for simulations that either include **(b)** MAN9 (cf. Figure 1b) or **(c)** FA2 glycans (cf. Figure 1c) on ACE2. The contact probabilities for both glycan systems (b & c) are virtually the same as panel (a), demonstrating that the RBD-ACE2 contact signature is robust. **(d)** Distributions of Cα-rmsd of RBD after least square fitting of ACE2, calculated for non-glycan and glycan simulations. **(e)** Structural representation of the RBD-ACE2 binding interface highlighting the residues that form persistent contacts (cyan in ACE2 and ice blue in RBD). Persistent contacts are those that form with at least 60% probability (see Table S1).

The contact maps for the non-glycan (Figure 2a), MAN9-included (Figure 2b), and FA2-included (Figure 2c) simulations are highly similar among each other, suggesting that RBD-ACE2 protein contact formation is virtually independent of the presence of glycans. To describe how glycans may affect the movement of the RBD domain relative to ACE2, we calculated the root mean square deviation (rmsd) of the C-alpha atoms of RBD after alignment of ACE2, as a function of time. From this, we calculated the distribution of rmsd values for each simulation set (Figure 2d). The distributions for the non-glycan and glycan-included simulations resemble each other closely, indicating that the glycans do not affect the orientation of RBD relative to ACE2. Specifically, in all simulations, the RBD fluctuates about a native-like basin that is located at small rmsd values (rmsd ≈ 0.5 nm, Figure 2d). Small rmsd values suggest that this ensemble corresponds to the RBD-ACE2 configuration used as reference for the rmsd calculations. Here, the structure by Ref. [5] (PDB ID: 6M0J, Figure 1a) was used as reference. Together, the unperturbed results for contact formation (Figure 2a-c) and rmsd distributions (Figure 2d) demonstrate that features including interactions and relative orientations of RBD-ACE2 are robust to the presence of glycans.

### Quantifying the relative significance of RBD-ACE2 contacts

To further characterize the RBD-ACE2 interaction profile, we defined persistent contacts as those that occur in at least 60% of the sampled configurations. Our simulations reveal that there are 25 RBD-ACE2 contacts that form persistently, which may be the key interactions in the binding of the complex (Figure 2e and Table S1). The majority of persistent contacts involve residues in the N-terminal helix of ACE2, which interact with the receptor-binding motif (RBM) of RBD (Figure 2e). Representative RBD-ACE2 interactions that occur most frequently in the simulations include: Y453-H34, F456-T27, N487-Y83, Y489-F28, N501-K353, G502-K353, and Y505-K353 (first and second labels denote RBD and ACE2 residues, respectively). These pairs form with at least 90% probability, suggesting they may play a particularly prominent role in the interplay between the virus’ RBD and human receptor (see Table S1 for a complete list of all 25 persistent contacts). In fact, of these seven RBD residues, two are sites where the common mutations Y453F and N501Y occur, and the remaining five are highly conserved among pandemic strains. Specifically, Y453F is a mutation that originates from minks in Denmark and has been found in over 1, 400 out of ~ 314, 000 sequences recorded globally (i.e. 0.45% global frequency). The N501Y RBD mutation, which appears mostly in the B.1.1.7 strain (or UK strain), has been found in over 18,000 out of ~ 314,000 sequences (i.e. 5.7% global frequency). The B.1.1.7 strain, first detected in the UK in September 2020, has become a dominant variant of the virus and is believed to be more transmissible. In the next section further below, we apply our contact statistics, as shown in Figure 2a, to assess how the N501Y mutation in B.1.1.7 may facilitate virus-receptor binding.

To discuss our results in view of recent experimental structures, we used the contact definition described above and identified all RBD-ACE2 contacts that are present in PDB IDs: 6M0J [5], 6M17 [6], and 6VW1 [7]. We find that these structures together implicate a total of 42 contacts. Since experimental models are based on average coordinates, it is not clear to what extent these 42 interactions statistically form or break in solution. Our simulations reveal that the experimentally-derived contacts can have substantially diverse probabilities of forming, ranging between *P* = 0.04-0.98 (Table S1). Notably, all 25 persistent contacts captured by the simulations represent a subset of those 42 experimentally-reported interactions. Thus, our analysis demonstrates that while recent structures have revealed the possible interactions, the present calculations quantify their relative involvement, which can aid in estimating the potential impact of mutations. We note that previous shorter simulations (~ 5 μs) [8] implicated a significantly higher number of persistent contacts than identified by our long simulations (~ 225μs). Such discrepancy highlights the importance of extensive sampling as performed here to obtain accurate statistics. Using large sets of simulation data, we have provided a more precise evaluation of RBD-ACE2 contacts, and have additionally shown that these interactions are not perturbed by the presence of MAN9 or FA2 glycans. Below, we demonstrate how our contact data may be applied to evaluate potential con-sequences of specific mutations, such as N501Y in the recently prevalent B.1.1.7 strain from the UK.

### Simulated interactions help assess the effects of mutations, such as N501Y in RBD

The detailed contact interactions, as revealed by our simulations (Figure 2a), provide a basis that can help evaluate the effects of mutations. In recent months, the new variant of SARS-CoV-2 B.1.1.7 (or UK strain), carrying the mutation N501Y in the RBD (Figure 3), reportedly accounts for over 60% of new COVID-19 cases in South East England. This N501Y variant is associated with increased affinity to host receptor [3], but the molecular factors responsible for this is unclear. Since the N501Y mutation site is at the RBD-ACE2 binding interface (Figure 3a), we will examine how this mutation may modulate the virus-host interaction, a factor that can determine infectivity. For this, we performed in-silico mutagenesis of N501Y in the RBD-ACE2 complex, using the FoldX software [19]. Utilizing the interaction statistics from Figure 2a as basis, together with a structural and chemical-physical analysis, we argue below how the new N501Y strain facilitates the binding of RBD to ACE2, which may contribute to enhanced infectivity.

**FIG. 3.**
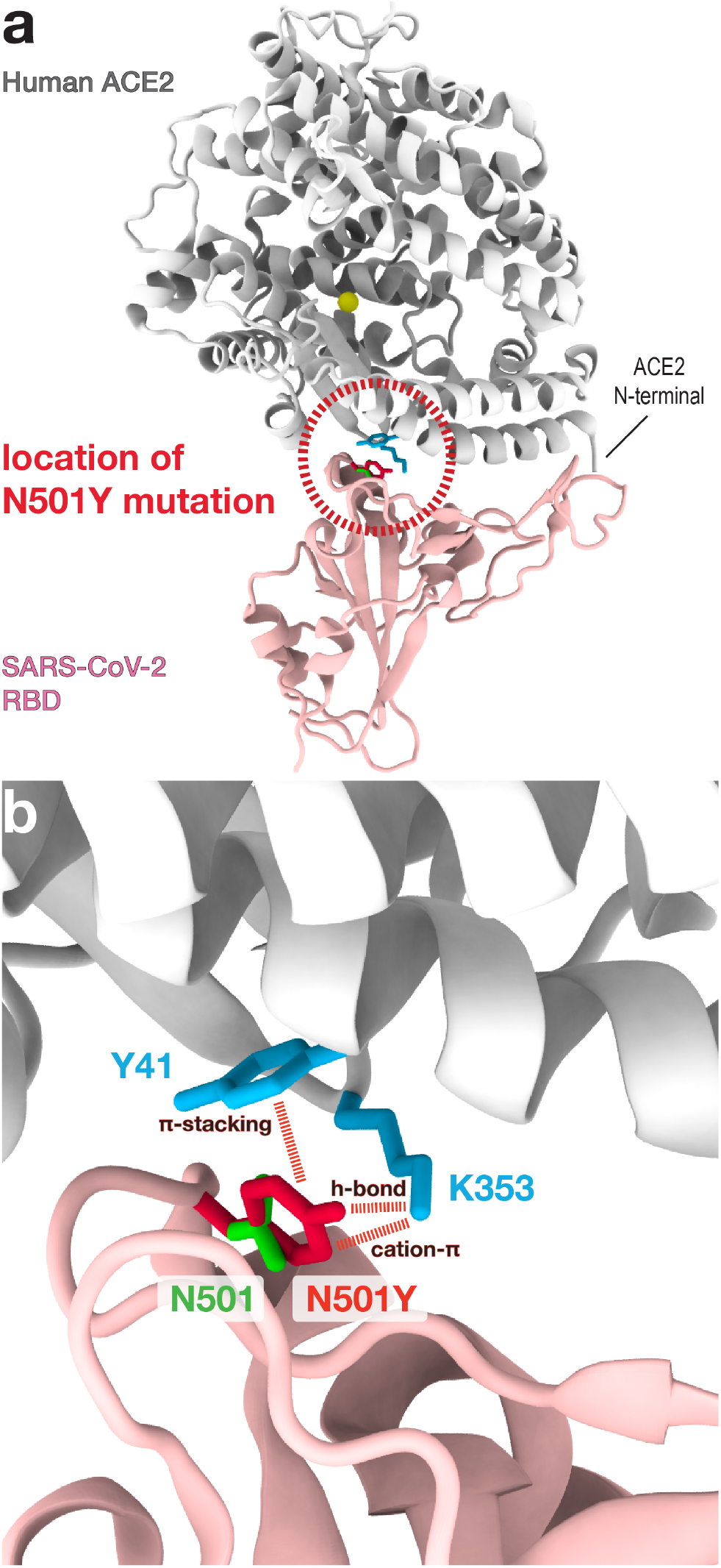
N501Y mutation introduces additional stabilizing interactions between RBD and ACE2. **(a)** Dashed circle indicates the location of the N501Y mutation site in RBD, which is at the binding interface with ACE2. **(b)** Zoomed-in view of N501Y location (i.e. dashed circle region of panel a). The wild type residue N501 of RBD is shown in green, and the mutation N501Y in red. The simulations show that N501 forms persistent contacts with Y41 and K353 (cyan) of ACE2 (cf. Figure 2a). Based on this, we assess how the N501Y mutation may alter the interactions with these ACE2 residues. As shown by dashed lines and their labels, N501Y would introduce additional stabilizing interactions with Y41 and K353 of ACE2, which will increase the binding affinity between RBD and ACE2.

To describe the con-sequences of N501Y, we first identify all ACE2 residues that frequently interact with N501, the mutation site in RBD. From the simulations, N501 forms contacts with Y41 and K353 of host ACE2 in 74% and 93% of the sampled configurations, respectively (see Figure 2a and Table S1). The significant interactions provide reason to specifically focus on these two host residues and assess how their contact formation with the N501Y mutation may alter stability (Figure 3b). For this, we note that N501Y would enable the formation of stabilizing cation-π interaction with K353 of ACE2. In addition, the longer side chain of N501Y (relative to N501) may facilitate the intermolecular hydrogen-bonding between the OH group of N501Y (tyrosine) and ACE2 residues, including K353 (Figure 3b). It has been shown that hydrogen bonds by tyrosine OH groups can be a significant contributor to stability [20]. Hence, a tyrosine (N501Y) side chain that allows for more favorable hydrogen bonds between molecules would further facilitate binding. Moreover, the N501Y mutation introduces additional stabilizing contributions: i.e. the aromaticaromatic interaction between N501Y and Y41 (i.e. π-stacking; Figure 3b). Notably, N501Y would lead to enhanced hydrophobic effects in the inside of the binding surface. This mutation would therefore create a more protein core-like environment, which stabilizes the RBD-ACE2 coupling. Together, our analysis demonstrates how N501Y implicates numerous energetic factors whose combined effect likely facilitates the binding of the mutated virus to host cells. This interpretation is consistent with the mutational experiments by Starr et al. [3] that have demonstrated enhanced RBD-ACE2 affinity through N501Y.

### RBD forms contacts with Asn90 glycan of ACE2 more frequently than other glycans

Some glycans on ACE2 are located near the RBD-ACE2 interface (Figures 1b,c). While we have shown that glycans do not alter the contact signature of ACE2 and RBD, it is unclear how and to what extent glycans of the host may interact with viral RBD. Understanding precisely which glycans form close contacts with RBD will provide a more comprehensive view of the factors determining affinity and may suggest additional strategies to modulate binding.

To describe RBD-glycan interaction, we calculated the probabilities of RBD residues forming contacts with any glycan that is attached to ACE2 (Figure 4). This analysis was performed for the simulation sets that either included MAN9 (cf. Figure 1b) or FA2 glycans (cf. Figure 1c) on ACE2. In both MAN9 and FA2 simulations (Figures 4a,b), ACE2 glycans on Asn53, Asn90, Asn103, and Asn322 form contacts with the RBD. Of these four glycans, the ones on Asn53, Asn103, and Asn322 have lower contact probabilities (i.e. P ≤ 30%), suggesting that their interaction with RBD is more transient despite their proximity to the binding surface. In stark contrast, the remaining glycan on Asn90 of ACE2 contacts the RBD with distinctly higher probabilities (P ≥ 70%), for both MAN9 and FA2 species (Figures 4a,b and S2). Specifically, this Asn90 glycan frequently forms contacts with residues of the RBD ranging approximately from 403-417 (Figures 4a-c). Notably, this RBD region does not form any contacts with ACE2 during simulation, regardless of whether glycans were present or absent (Figure 2a-c). Since RBD-ACE2 contacts are independent of the presence of glycans (as discussed earlier), this result indicates that contact formation between RBD and the Asn90 glycan of ACE2 does not compete with the protein-protein interactions between RBD and ACE2.

**FIG. 4.**
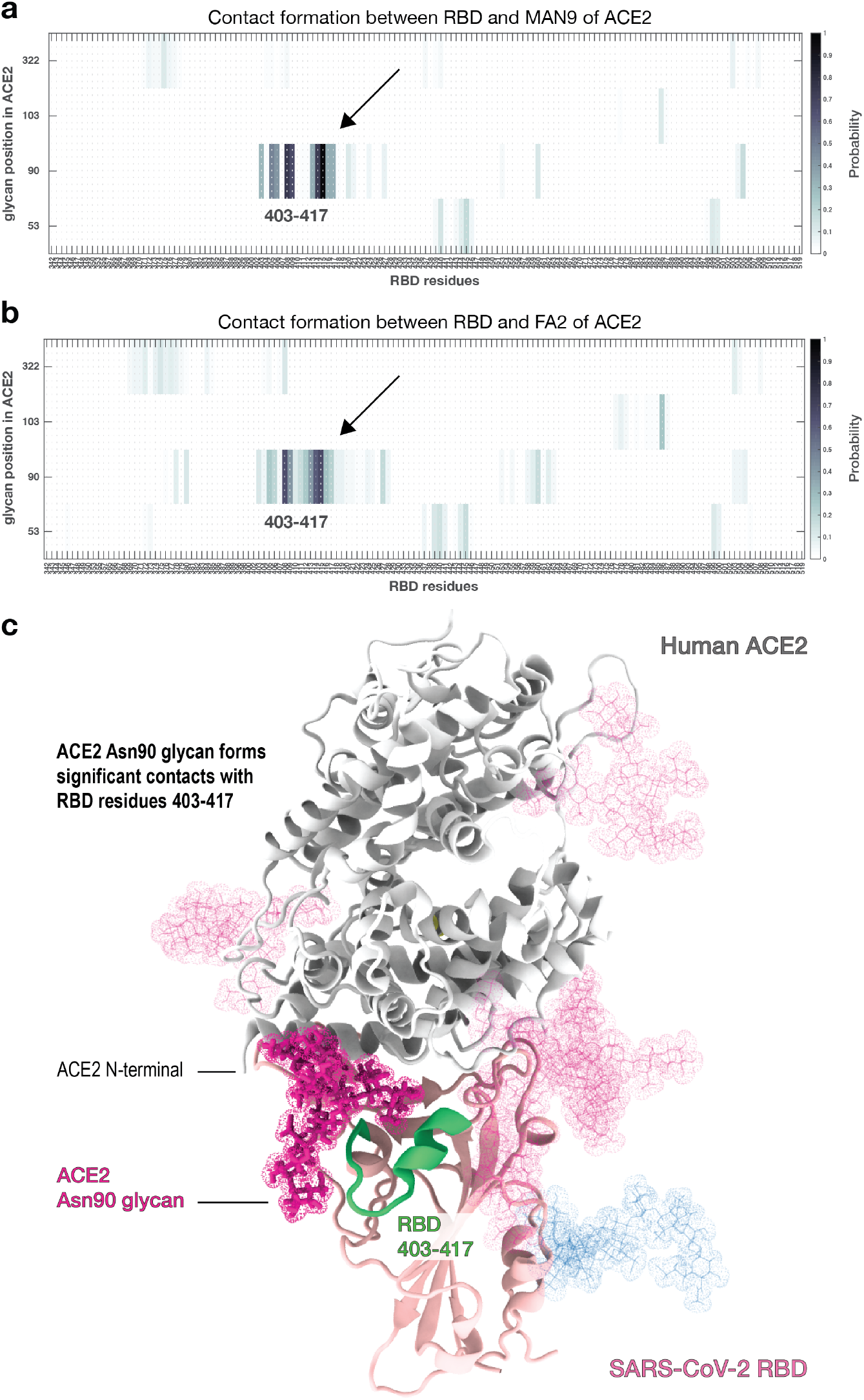
Only the glycan on Asn90 of ACE2 forms significant contacts with RBD. **(a)** Probability of contact formation between RBD and MAN9 glycans of ACE2. **(b)** Contact probability between RBD and FA2 glycans of ACE2. Both (a) and (b) show that only the contacts involving Asn90 glycan occur with higher probabilities (see region indicated by arrow). RBD residues that form these protein-glycan interactions are roughly between 403–417. This RBD region does not form contacts with ACE2 in the non-glycan or glycan-included simulations (cf. Figure 2a-c). This suggests that the glycans do not compete with protein-protein interactions at the RBD-ACE2 interface. **(c)** Structural description of the ACE Asn90 glycan (magenta) and RDB residues 403–417 (green), forming glycan-protein interactions.

The prominent role of the Asn90 glycan as elucidated by our long, unrestrained simulations is consistent with previous atomic force microscopy (AFM) measurements and steered molecular dynamics (SMD) simulations (~ 270 ns) [16]. In that study, AFM and SMD analyses suggest that the Asn90 glycan can affect the association and disassociation of RBD and ACE2. As shown in Figure 4, the frequent contacts between Asn90 glycan and RBD implies that there are significant steric effects associated with this glycan. Since RBD-ACE2 interface interactions are not perturbed by glycans (as discussed above), the sterics of Asn90 glycan would hinder the unbinding of RBD, hence stabilizing the RBD-ACE2 complex. This result helps explain why, in AFM unbinding experiments [16], separating RBD and ACE2 requires higher forces when ACE2 glycans are not removed. For the case when ACE2 is not bound to RBD, the excluded volume of the Asn90 glycan is likely to interfere with the binding interface, thereby impeding the association of RBD. Consistent with this interpretation, mutation studies have shown that removal of the Asn90 glycan leads to increased RBD binding events [17, 18]. Together, our simulations provide a mechanistic basis for how the sterics of Asn90 glycan can impede the unbinding and binding of RBD.

In previous SMD simulations [16], which reported on glycan-RBD interactions, all glycans of ACE2 were modeled as N-glycan core pentasaccharide, which is a minimum structure for all N-glycans. Thus, the results from Ref. [16] may not apply to specific glycan types that have distinct structures or chemical-physical properties. Here, we specifically included MAN9 or FA2 glycans in ACE2 in our simulations and have shown that they are associated with comparable glycan-RBD contact interactions (Figure 4a,b). While the contacts made with RBD may be similar for MAN9 and FA2, different chemical-physical properties of the glycans can lead to distinct binding energetics of the RBD-ACE2 complex, as will be discussed in a later section of this manuscript.

### Interactions between Asn343 glycan of RBD and glycans of ACE2 may contribute to affinity

In addition to protein-protein and protein-glycan interactions, glycan-glycan interactions can also be a contributor to RBD-ACE2 binding, and thus, infectivity. On the Spike RBD, we modeled the glycan at Asn343 as a complex FA2 glycoform, in accordance with previous experimental studies [9, 21]. In our simulations, the Asn343 glycan of RBD forms contacts with the Asn53 and Asn322 glycans of ACE2 in 20% and 40% of the sampled configurations (regardless of ACE2 glycan type, see Figure S3), suggesting that these interactions may be a determinant of the stability of the complex. Consistent with this notion, SARS-CoV-2 pseudovirus essays have shown that removal of the RBD-Asn343 glycan leads to a 20-fold decrease in infectivity [11]. The functional relevance of the RBD-Asn343 glycan may be reflected by the fact that this glycan is highly conserved in current GISAID SARS-CoV-2 sequences. Specifically, it is lost in only 3 out of 313, 826 sequences, according to cov.lanl.gov Spike alignment (as of January 16, 2021). Interestingly, the RBD-Asn343 glycan forms more frequent contacts only with the two glycans of ACE2 mentioned above (i.e. the ones at Asn53 and Asn322), but does not form significant interactions with any protein residues of ACE2 (less than 10% contact probabilities, see Figure S3). This suggests that the RBD-Asn343 glycan may contribute to infectivity, as implicated by [11], predominantly through interactions with the Asn53 and Asn322 glycans of ACE2 (Figure S3).

### RBD-ACE2 binding energetics depend on glycan type

To elucidate how the energetics of RBD-ACE2 binding may be affected by MAN9 or FA2 glycans, we applied the Molecular Mechanics Poisson-Boltzmann Surface Area (MM-PBSA) approach [22, 23]. We employ this end-state free-energy calculation to estimate the relative changes in binding of RBD and ACE2, with and without glycans. In the present MM-PBSA calculations, the binding energy of a ligand-receptor complex (RBD and ACE2) is approximated by changes in molecular mechanics and solvation energies. Here, the sum of the two contributions is referred to as MM-PBSA energy, which is composed of enthalpic terms and an approximation for the solvent entropy (see SI Methods for details). To assess the free energy of binding (i.e. affinity), one needs to further determine changes in conformational entropies. Separately from the MM-PBSA energy calculation, the configurational entropy was estimated using a quasi-harmonic approach. We found that the relative changes in entropy are similar for the MAN9 and FA2 simulations, and therefore, did not include them in the binding energy calculations (see SI Methods and Figure S4a). Hence, the MM-PBSA energies discussed below can be used to approximate relative changes in binding energetics, allowing us to infer how each glycan type can influence RBD-ACE2 affinity.

To dissect the effect of different glycan types on RBD-ACE2 stability, we calculated MM-PBSA energies (i.e. stability) for the simulations where glycans were not included, and where MAN9 or FA2 glycans were bound to ACE2 (Figure 5a). For each simulation set, we used large numbers of representative snapshots for MM-PBSA analysis to ensure statistical convergence (see SI Methods for details). We find that the simulations with MAN9 glycans on ACE2 result in a 14.7% decrease in RBD-ACE2 binding stability, relative to the simulations without glycans (Figure 5a). In contrast, the trajectories with FA2 glycans on ACE2 lead to a 9.1% increase. While this MM-PBSA analysis does not report free energies, the distinct combinations of enthalpy and solvent entropy demonstrate that RBD-ACE2 affinity can be dependent on glycan species. Specifically, since the entropies associated with MAN9 and FA2 glycans are comparable (Figure S4a), our calculations suggest that FA2 glycans in ACE2 would have a stabilizing effect on the RBD-ACE2 complex, while the MAN9 type would lead to weaker binding affinity.

**FIG. 5.**
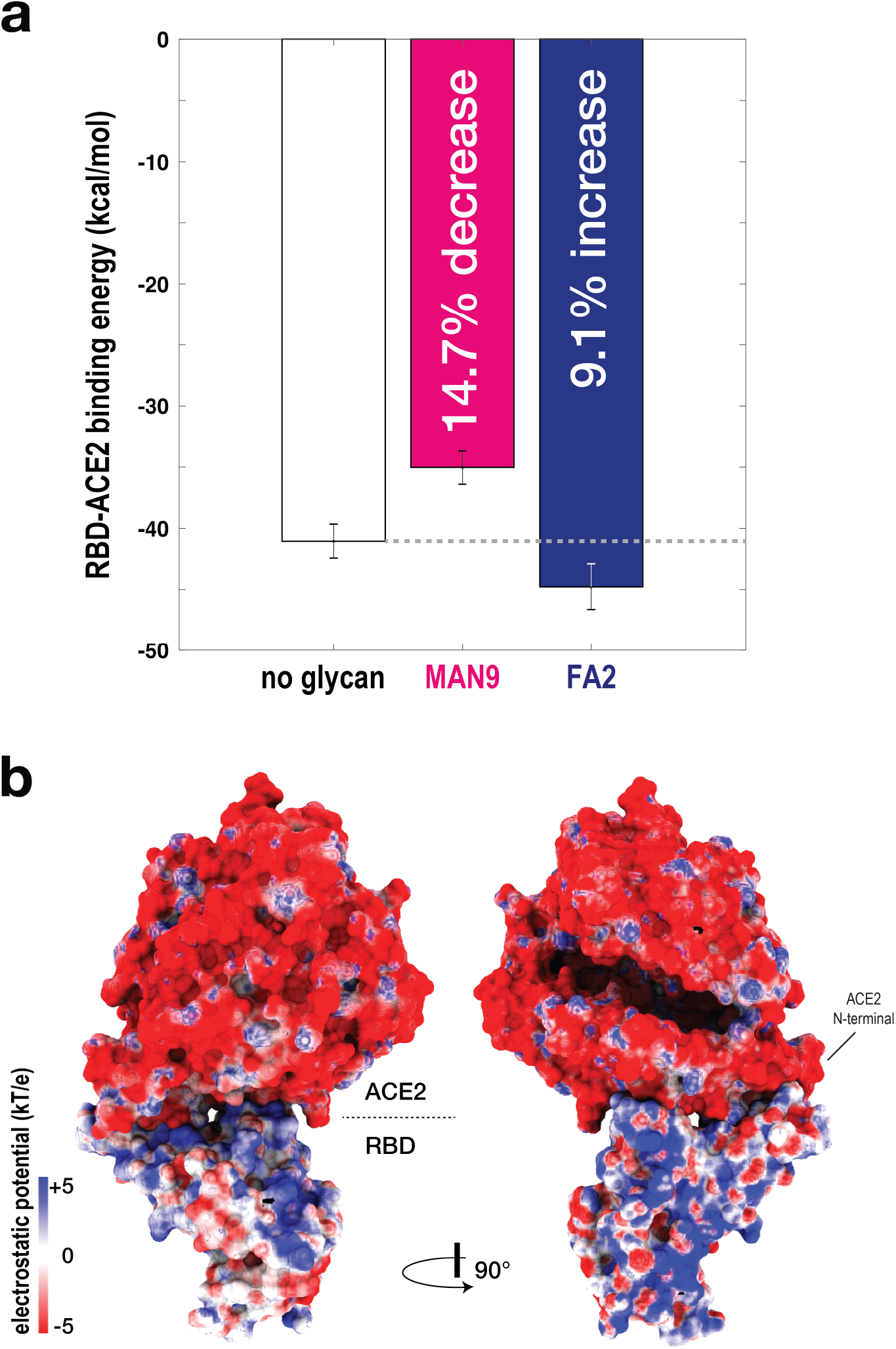
RBD-ACE2 binding affinity is glycan dependent. **(a)** To evaluate the binding energy between RBD-ACE2, the MM-PBSA approach was applied. Binding energy was calculated for simulations without glycans (white bar), and with MAN9 glycans (magenta bar) or FA2 glycans (blue bar) on ACE2. Simulations with MAN9 glycans on ACE2 are associated with a 14.7% decrease in stability, relative to the non-glycan simulations. In contrast, simulations with FA2 glycans on ACE2 result in a 9.1% increase in stability. **(b)** Electrostatic surface potential calculated for ACE2 and RBD. Two complementary views show that the ACE2 surface is overall negatively charged, while the surface of RBD is overall positive.

As shown earlier, of the six glycans that are on ACE2, only the one at Asn90 forms significant contacts with RBD, regardless of ACE2-glycan type (cf. Figures 4a,b and S2). Since only glycan Asn90 forms close contacts with RBD, one could ask if the binding energy perturbations (as presented in Figure 5a) resulted solely from the effects of glycan Asn90. To answer this, we removed glycan Asn90 from both trajectories where either MAN9 or FA2 glycans were on ACE2, and repeated the MM-PBSA analysis. When glycan Asn90 is excluded (Figure S4b), the MAN9 trajectory leads to a 9.6% decrease in RBD-ACE2 binding stability, relative to the simulations without glycans (contrast this to the 14.7% decrease for when glycan Asn90 is present, cf. Figure 5a). For the FA2 simulations, excluding glycan Asn90 results in an increase of 3.6% in stabilizing energy (versus the 9.1% increase for when glycan Asn90 is included, cf. Figure 5a). This comparison demonstrates that while glycan Asn90 is a dominant contributor to the observed affinity perturbations, it is not the sole contributor out of the six ACE2 glycans. Since the remaining glycans on ACE2 lead to appreciable stability changes (Figure S4b) while not forming close contacts with RBD (Figures 4a,b and S2), it suggests that glycans affect RBD-ACE2 affinity through long-range electrostatic interactions, as discussed in more detail below.

### Electrostatic effects of different glycan types lead to distinct RBD-ACE2 binding energetics

To identify which energetic contributions are responsible for the distinct affinities as shown in Figure 5a, we dissected individual components of the MM-PBSA energy for the simulations without glycans, and those with MAN9 or FA2 glycans in ACE2. Relative to the non-glycan simulations, the ones with MAN9 or FA2 glycans in ACE2 each lead to a similar increase in stabilizing van der Waals energetics of roughly 30% (Figure S4c). In terms of solvation energy (summation of polar and nonpolar), the MAN9 and FA2 simulations are associated with greater stabilizing contributions by 35% and 31%, compared to the non-glycan case. Regarding electrostatic energy, the trajectories with MAN9 or FA2 glycans in ACE2 result in decrease of stabilizing contributions by 42% or 32%, relative to the value for non-glycan simulations. The summation of these energetic components constitute the MM-PBSA results shown in Figure 5a. Our energy partitioning demonstrates that electrostatic contributions, specific to each glycan type, are mainly responsible for the observed variation in binding affinity of RBD-ACE2.

As just discussed above, electrostatic effects are the dominant factor for the observed difference in MM-PBSA properties associated with MAN9 and FA2 glycans. Distinct electrostatic features can stem from the charged sialic acids in FA2 and the characteristic solubility of each glycan type. It has been suggested that the ACE2 binding surface is overall negatively charged, while the corresponding RBD interface is positive [24]. Here, we further determined that ACE2 and RBD surface-potential distributions are also mainly negative and positive, respectively, for regions well beyond the binding interface (Figure 5b). Hence, interaction of the positive RBD with the negative sialated tips of the FA2 glycans on ACE2 would lead to stabilizing MM-PBSA contributions, as compared to MAN9 glycans that do not have these sialated tips. Since the RBD-ACE2 contact map is independent of the presence of glycans, the observed MM-PBSA effects should be mainly the result of long-range electrostatic interactions of the sugars that are flexible and sample wide regions around the RBD. These findings are in agreement with numerous studies that have identified sialic-acid interaction sites away from the binding interface, which may be critical for virus-host binding [25, 26].

## DISCUSSION

To develop effective strategies for treating COVID-19, one requires thorough insights into the factors that determine the binding between viral RBD and human ACE2. To this end, recent structural analyses [5-7] and mutational experiments [3, 4] have alluded to possible RBD-ACE2 interactions, as well as the role that glycans may play during this process. To extend these initial findings, one must obtain a detailed mechanistic understanding of the dynamics associated with numerous contributors to binding. For this purpose, we have performed long all-atom simulations of the RBD-ACE2 complex in explicit solvent. To dissect each contributor, we simulated separate models where different glycan types are bound or excluded (Figure 1). Interestingly, the simulations show that MAN9 or FA2 glycans have virtually no effect on the interface contacts between RBD and ACE2 (Figures 2a-c), nor the fluctuations of RBD relative to ACE2 (Figure 2d). Our statistical evaluation of contacts has identified which RBD-ACE2 interface residues are most likely to form interactions, thereby pinpointing the critical sites for binding. This analysis is based on the most extensive RBD-ACE2 simulations to date, comprising over 225 μs, which is one to two orders of magnitude longer than previous simulations. Hence, our results represent more accurate statistics that may be used as reference for interpreting experimental measurements, or gauging the impact of specific mutations. As an example of application, we have used the simulations as guide to evaluate how the novel N501Y mutation may modulate the affinity between virus and host (Figure 3). Specifically, our analysis suggests that N501Y leads to additional stabilizing interactions (i.e. with Y41 and K353 of host ACE2, see Figure 3b) that can facilitate virus binding.

The present simulations suggest that excluded volume effects of the Asn90 glycan in receptor ACE2 may determine the binding and unbinding of viral RBD (Figure 4). Precise characterization of glycan structure and function in experiments has not been possible so far, because glycan structures are highly complex, heterogeneous, and flexible [27, 28]. All currently available SARS-CoV-2 Spike and human ACE2 structures only include glycans that are partially resolved (i.e. up to the minimal stem conformation). Furthermore, these incomplete glycan structures represent only a very limited conformational ensemble, since the flexible glycans can sample extremely large conformational spaces. Therefore, to accurately describe the effect of glycans, the simulations must capture sufficiently large number of configurations. To account for this, we have generated 40 long simulations with MAN9 or FA2 glycans, where each run was initiated using a distinctive starting configuration (see Methods for details). The simulated trajectories have revealed that the glycan on Asn90 of receptor ACE2 forms more significant contacts with viral RBD than any other ACE2 glycan. This suggests that steric effects from Asn90 glycan may impede the unbinding of RBD, thus stabilizing the RBD-ACE2 complex. These results are consistent with recent atomic force microscopy (AFM) experiments, where separating RBD and ACE2 requires higher forces when the Asn90 glycan is present [16]. On the other hand, when ACE2 is not in complex with RBD, the significant steric presence of Asn90 glycan indicates that it would obstruct the binding site, thereby hindering the RBD from associating with ACE2. This notion agrees with mutational experiments demonstrating that removal of the Asn90 glycan leads to increased RBD binding events [17, 18]. Here, by uncovering the prominent role of Asn90 glycan in forming contacts with RBD, our simulations indicate how Asn90-glycan sterics may impede the binding, as well as the unbinding of RBD.

With increasing focus on the use of soluble extracellular domains of ACE2 as decoy inhibitors, it is critical to understand how specific glycan types on ACE2 can affect the binding energetics of RBD-ACE2. Our calculations suggest that distinct electrostatic features of different glycan types can be decisive for binding affinity (Figure 5). Specifically, we have shown that stabilizing energetics arise from interaction between the overall positively charged RBD and the negative sialated tips of complex FA2 glycans on ACE2. These observations are in line with experiments that reported stabilizing binding sites for sialic acids on Spike proteins in various SARS and MERS strains [25, 29]. Sialation has also been known to improve thermal stability and solubility [30, 31], as well as better recognition of “self” versus “non-self” by Siglecs [32], which are all favored features for effective immunogens. Together, our simulations, along with these supporting experiments, help establish how glycosylation variability on ACE2 can contribute to the slight discrepancies in binding affinities of SARS-CoV-2 for host receptor [1, 5, 7, 33, 34], possibly explaining the broad range of host-immune responses in the human population [35].

The present study elucidates the numerous contributors to the interplay between viral RBD and human ACE2, including protein-protein interactions, and the effect of specific glycan types on the binding of the RBD-ACE2 complex. The simulations have provided evidence that the stability of RBD-ACE2 is dependent on which glycan type is bound to the host receptor. Remarkably, ACE2 glycans can affect virus binding affinity through electrostatic effects, while not perturbing the physical contacts that are formed between virus and host. Of the numerous glycans on ACE2, the calculations have revealed that glycan Asn90 is a dominant contributor to affinity perturbations, and may play a critical role in the binding and unbinding events of RBD. Together, our results provide a theoretical framework that may be used to design more precise experiments that aim at regulating the infectivity of SARS-CoV-2.

## METHODS

To elucidate the interaction between RBD and ACE2, we performed all-atom explicit-solvent molecular dynamics (MD) simulations of the RBD-ACE2 complex (PDB ID: 6M0J [5]; Figure 1a). In all models used for simulations, the SARS-CoV-2 RBD domain is defined by residues T333-G526, and the ACE2 receptor by S19-D615. To avoid artificial charges at the protein ends, we introduced N-terminal acetylated and C-terminal N-methylamide capping groups. The ACE2 structure contains a zinc ion that is coordinated by H374, H378, E402, and one water molecule [5, 36]. Zinc coordination plays a critical role in maintaining the structural integrity and stability of a protein [37]. To properly account for this in the simulations, we introduced bonded terms between the zinc ion and H374, H378, E402 in ACE2. Equilibration values for distances and angles of these bonded terms were defined by the values found in the RBD-ACE2 configuration of PDB ID: 6M0J [5]. The strength of the zinc interactions were set at values that are equivalent to covalent bond interactions, as described in the CHARMM36m forcefield [38] applied for the present simulations. Note that we determined these force parameters for our previous simulations of C-Raf CRD (unpublished data). In these C-Raf simulations, the parameters used here led to structural fluctuations of the zinc site, whose scale at 310 K is consistent with the NMR ensemble from Ref. [39].

To dissect the role of distinctive glycan species in the interaction of RBD and ACE2, we performed additional separate sets of simulations where (a) MAN9 and FA2 glycans are attached to ACE2 and RBD, respectively (Figure 1b, S1) and (b) FA2 glycans are on both ACE2 and RBD (Figure 1c). Generally, glycans have complex structures and are highly flexible, allowing them to sample very broad conformational spaces. Accordingly, simulations may not accurately capture the effect of glycans if these calculations depend on the choice of initial configuration. To prevent this artifact, we prepared 40 complementary initial configurations for the MAN9 and FA2 simulations (i.e. 20 for each simulation set). Each glycan initial structure was modeled based on a simulated RBD-ACE2 configuration from preliminary trajectories (see SI Methods for details on glycan modeling). By initiating the glycan simulations from many distinctive configurations, we capture larger conformational ensembles that can more accurately partition the contribution of each glycan type.

### Simulation Details

All-atom explicit-solvent simulations were performed with the AMBER 16 software package [40]. The CHARMM36m forcefield [38] and TIP3P water model [41] were used. Each configuration was solvated and centered in a cubic box. The size of the cubic box was chosen to create at least 15 Å padding on each side along the largest atom-atom distance of the molecule. Each system was neutralized with an excess of 150 mM KCL. Energy minimization was performed using the steepest descent algorithm. Equilibration simulations were first carried out under the constant number-volume-temperature (NVT) ensemble for 2 ns, and then under the constant numberpressure-temperature (NPT) ensemble for 10 ns. During both equilibration stages, harmonic position restraints were imposed on all non-hydrogen atoms of the molecule. Constant temperature was maintained at 310 K using velocity Langevin dynamics [42], with a relaxation time of 1 ps. Constant isotropic pressure of 1 bar was achieved by employing the Berendsen barostat [43], with a relaxation time of 4 ps and compressibility of 4.5 × 10^-5^ bar^−1^. Covalent bond lengths were constrained with the SHAKE algorithm [44]. Van der Waals interactions were evaluated using a cutoff where forces smoothly decay to zero between 1.0-1.2 nm. Coulomb interactions were computed using the particle-mesh Ewald (PME) method [45], with Fourier grid spacing of 0.08-0.10nm and fourth order interpolation. Unrestrained production simulations were performed in the NPT ensemble, with an integration time step of 4 fs, which was enabled through hydrogen mass repartitioning [46].

For the non-glycan RBD-ACE2 model (Figure 1a), 50 simulations were performed, with a total simulated time of 165 *μs* (i.e. each trajectory includes roughly 3.3 μs). For the complex with MAN9 glycans in ACE2 (Figure 1b), 20 simulations were performed for an aggregated time of over 30 μs (each replica of this set is about 1.5 μs long). Finally, the simulation set for the model with FA2 glycans in ACE2 (Figure 1c) contains 20 simulations that exceed 30 μs of accumulated time (each trajectory of this set is roughly 1.5 μs long).

## ADDITIONAL INFORMATION

**Supporting Information** accompanies this paper.

## ACKNOWLEDGMENTS

K.N. and S.G are partially supported by the DOE Office of Science through the National Virtual Biotechnology Laboratory, a consortium of DOE national laboratories focused on response to COVID-19, with funding provided by the Coronavirus CARES Act. S.C. is supported by the Center of Nonlinear Studies Postdoctoral Program. R.A.M. was supported by a Los Alamos National Laboratory (LANL) Director’s Postdoctoral Fellowship. B.K. and S.G. are partially supported by LANL LDRD project 20200706ER. This research used computational resources provided by the LANL Institutional Computing Program.

## AUTHOR CONTRIBUTIONS

K.N., S.C., R.A.M., B.K., and S.G. designed research. K.N. and S.C. performed simulations. S.C., K.N., and R.A.M prepared structural models. S.C., K.N., R.A.M., B.K., and S.G. analyzed data. K.N. and S.C. created figures and wrote the paper. K.N., S.C., R.A.M., B.K., and S.G. edited the paper. B.K. and S.G. obtained funding.

## Notes

The authors declare no competing interest.

## Supporting Information

### SUPPORTING METHODS

#### Choice of glycosylation and glycan modeling

The ACE2 receptor has 7 potential N-glycosylation sites (PNGS), N53, N90, N103, N322, N432, N546, and N690. Of these, the first six glycans are present in the ACE2 residue range used in our simulations (i.e. S19-D615). It remains to be seen exactly how different glycan types at these sites affect ACE2 association with the viral RBD. It is known that post-translational glycan modifications are strongly dependent on expression cell lines and their glycosylation enzyme repertoire [1,2]. Unfortunately, all currently available ACE2 studies were done using recombinant proteins expressed in non-native cells. This prevents a definite determination of native glycosylation pattern on the ACE2 receptor and their role in RBD binding. Literature suggests that DC-SIGN and L-SIGN lectins act as enhancer factors that facilitate ACE2 mediated virus infection [3]. These specifically recognize high-mannose glycans [4], indicating that at least those glycans on ACE2 interacting with these lectins occur in oligomannose form. On the other hand, Zhao et al. [5] had previously applied sequential exo-glycosidase digestion to identify mainly biantennary N-linked glycans with sialylation and core fucosylation. Recently, Shajahan et al. [6] performed site specific mass spectrometry analysis of human ACE2 to indicate predominantly complex type glycosylation, with 60% biantennary, 85% fucosylated, and about half of them as sialated structures. Moreover, negatively charged sialic acids extensively found on complex glycans have been reported to play critical roles in viral Spike interaction [7]. A thorough understanding of the effects of glycosylation is thus necessary.

Since each PNGS can have a varying distribution of glycan occupancies [6], we modeled two divergent forms of N-glycans on the different ACE2 sites, namely the unprocessed 9-mannose (MAN9) oligomer and the enzymatically processed fucosylated 2-antennae type complex glycan (FA2) with commonly expected 2-3 linked [6] sialic acid tips. Since FA2 glycan type has been shown to be the major glycoform at the Spike glycosylation site 343 [8, 9], this was selected as the glycan choice for RBD for all glycan-included simulations. 40 initial configurations were modeled for the MAN9 and FA2 simulations (i.e. 20 for each simulation set; cf. main text Figure 1b,c). These initial configurations were prepared based on different RBD-ACE2 configurations taken from preliminary glycan-free trajectories. Glycan structures were built at the PNGS, with random orientations, using the ALLOSMOD package [10] of MODELLER [11]. This was succeeded by short simulated annealing with the protein backbone restrained to relax the glycosylated systems at different conformations, with the CHARMM36m forcefield [12], following the glycoprotein modeling pipeline developed previously by our group [13, 14]. The CHARMM pdb and psf files were converted to AMBER format using the CHAMBER command available in the PARMED module of AMBERTOOLS 16 [15, 16]. Following the steps described, 20 different glycoprotein configurations were obtained for each of the MAN9 and FA2 glycosylated ACE2 systems, which were used for the 40 individual glycosylated trajectories of all-atom explicit-solvent simulations performed with the AMBER 16 software [15].

#### MM-PBSA calculations

The binding energy between RBD and ACE2 was approximated using the Molecular Mechanics Poisson-Boltzmann Surface Area (MM-PBSA) method [17, 18]. To apply this method, we used the MMPBSA.py script [19] within AMBERTOOLS 16. MM-PBSA estimates the binding energy (Δ*G*_bind_) from the molecular mechanical energy (Δ*E*_MM_), solvation free energy (Δ*G*_sol_) and conformational entropy *(ΔS*) as:

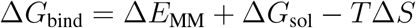

with

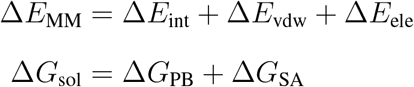

*T* is the temperature; Δ*E*_int_ is the internal energy from the sum of bond, angle, and dihedral terms; Δ*E*_vdw_ is the van der Waals energy; Δ*E*_ele_ is the electrostatic energy; Δ*G*_PB_ is the electrostatic solvation free energy computed by the Poisson-Boltzmann (PB) method [20]; and Δ*G*_SA_ is nonpolar solvation free energy proportional to solvent accessible surface area and cavitation terms. ΔGSA implicitly includes the solvent entropy approximation by virtue of parameterization [21, 22]. Because the PB calculations are computationally very costly, 4 sets of randomly selected 200 snapshots were used from the complete ensembles for these calculations in order to obtain robust sampling and standard errors.

MM-PBSA has been shown to perform reasonably well for protein-glycan systems in order to calculate relative affinity changes and their agreement with experimental values [23]. The conformational entropy change (ΔS) is not included in the present MM-PBSA analysis since entropy calculations are typically error-prone [24,25] and have convergence difficulties [26,27]. It has been shown that the inclusion of entropic terms from quasi-harmonic approximation provided no meaningful improvement in the agreement between the predicted and experimental energies, whereas other methods of conformational entropy calculations such as harmonic approximation entropies reduced the correlation [23]. Here, the quasi-harmonic conformational entropy was calculated from the eigenvectors of complete covariance matrix in GROMACS v5.1.2 [28] (Figure S4a). Since the relative changes in conformational entropy are similar between the MAN9 and FA2 simulations (Figure S4a), we did not included them in our binding energy calculations. Electrostatic potential calculation was performed using the Adaptive Poisson Boltzmann Solver [29].

### SUPPORTING FIGURES AND TABLES

**FIGURE S1.**
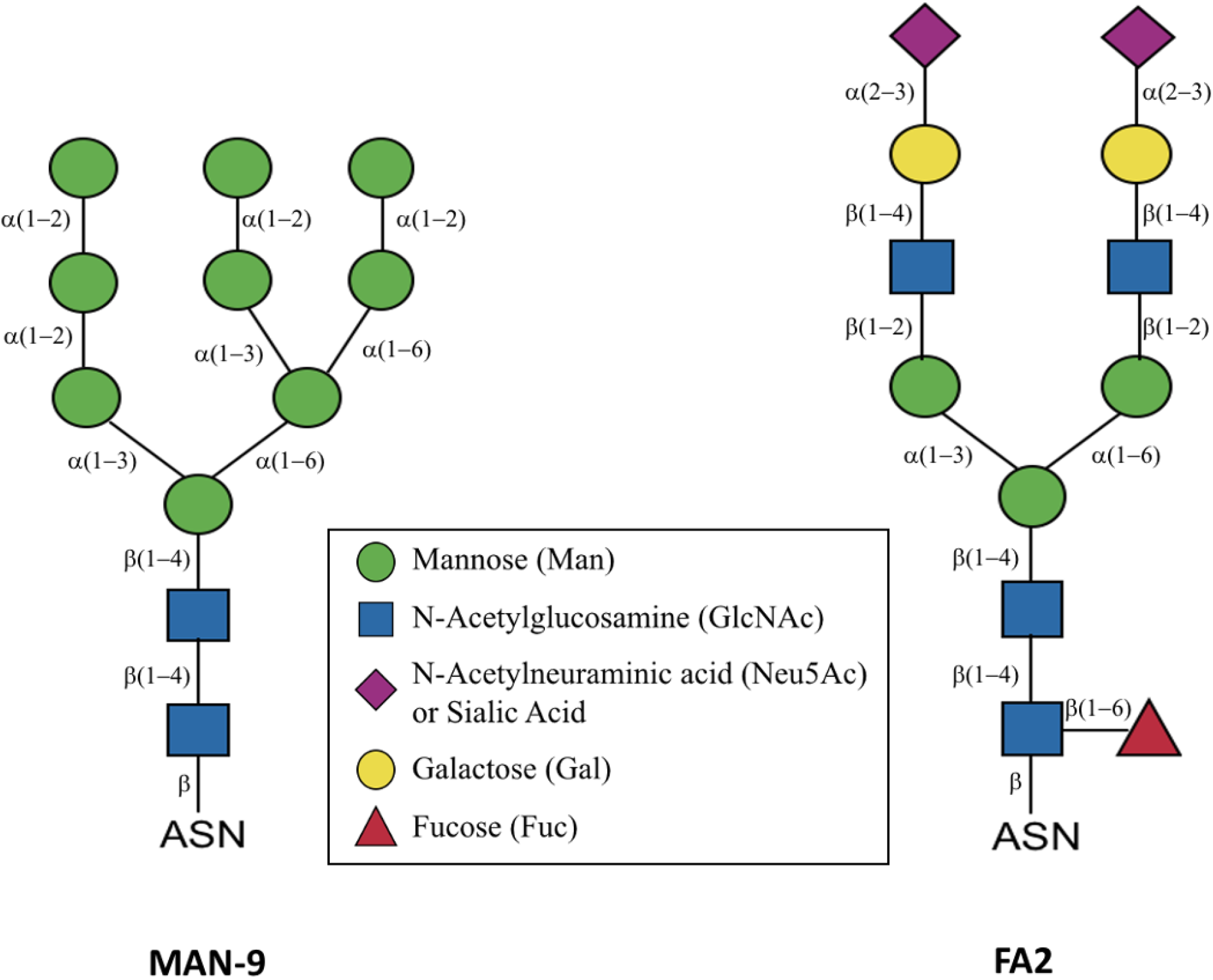
Schematic representation of N-glycans. Oligomannose (MAN9) and complex glycan (FA2) as used in the simulations, represented in Symbol Nomenclature for Glycans (SNFG) schematic. The inter-sugar connectivities are shown.

**FIGURE S2.**
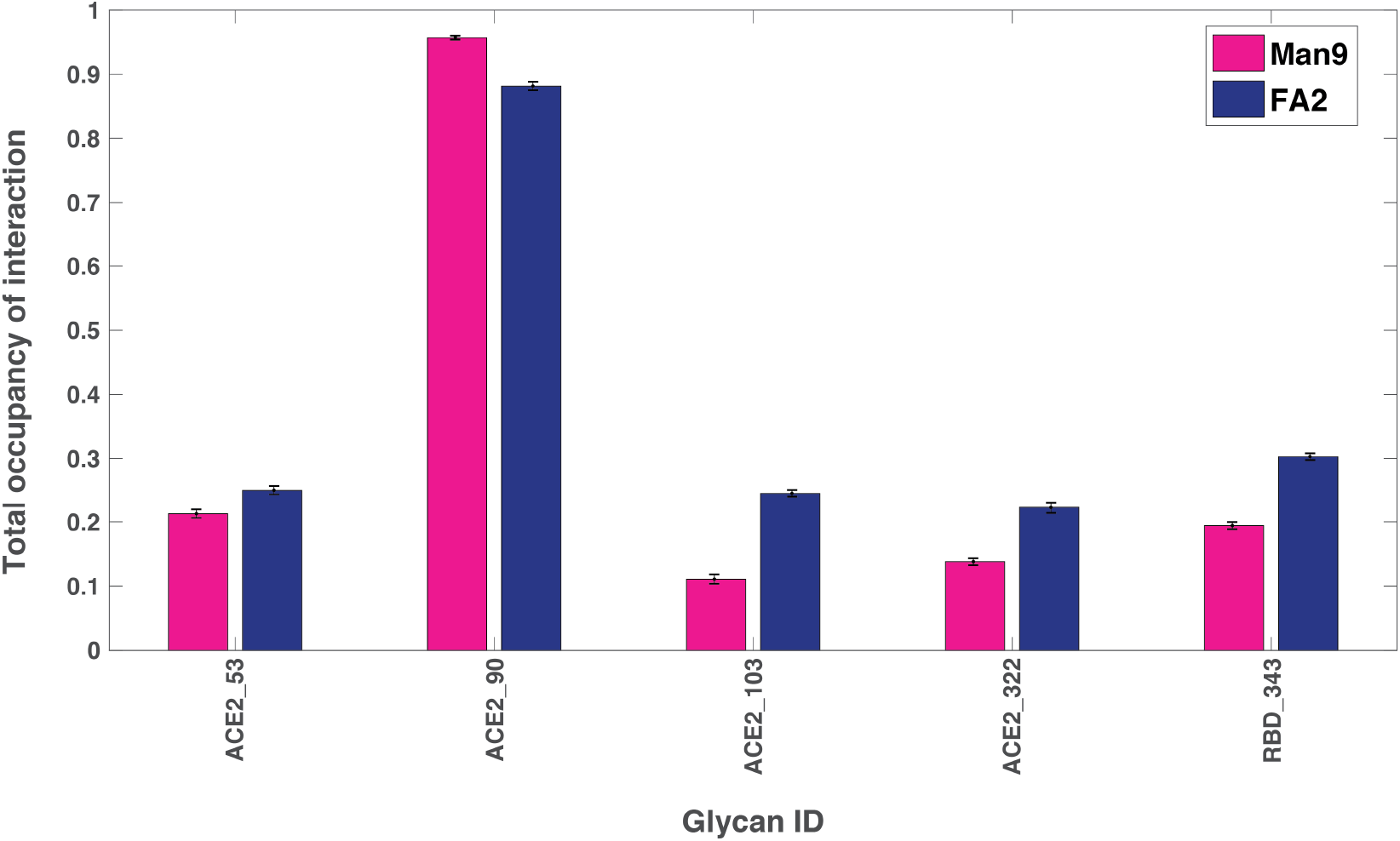
Total probability of glycan-protein contact formation for each glycan. Of all the glycans in ACE2, the one at position 90 contacts the RBD with distinctly higher probability (over 80%). The single RBD glycan at position 343 forms contacts with ACE2 in 20% (30%) of the sampled configurations for MAN9 (FA2). Standard deviations by bootstrapping over four sets from the total ensemble demonstrate convergence of glycan sampling.

**FIGURE S3.**
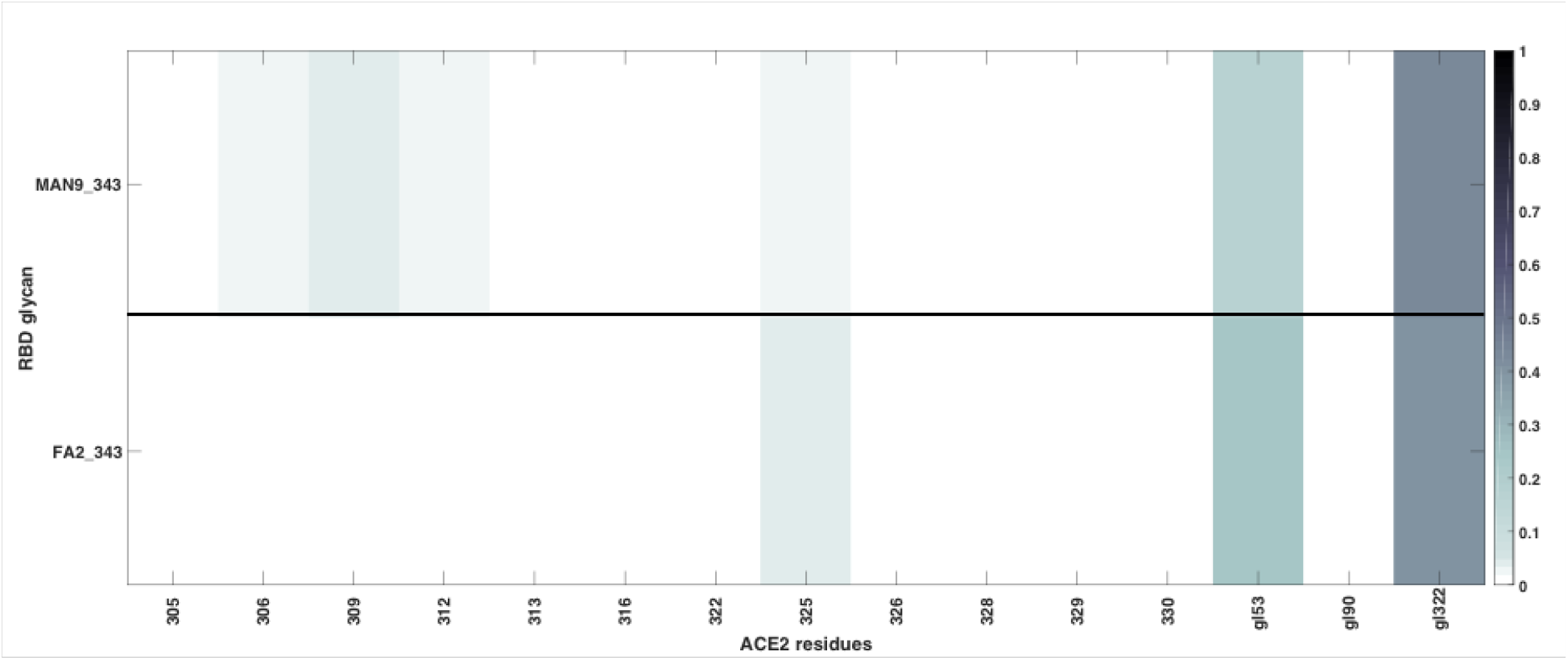
Interactions of RBD glycan 343 with ACE2 residues or ACE2 glycans. RBD glycan 343 forms more frequent interactions (over 20%) with glycans 53 and 322 of ACE2 (both for MAN9 and FA2). RBD glycan 343 does not form frequent contacts with ACE2 residues.

**FIGURE S4.**
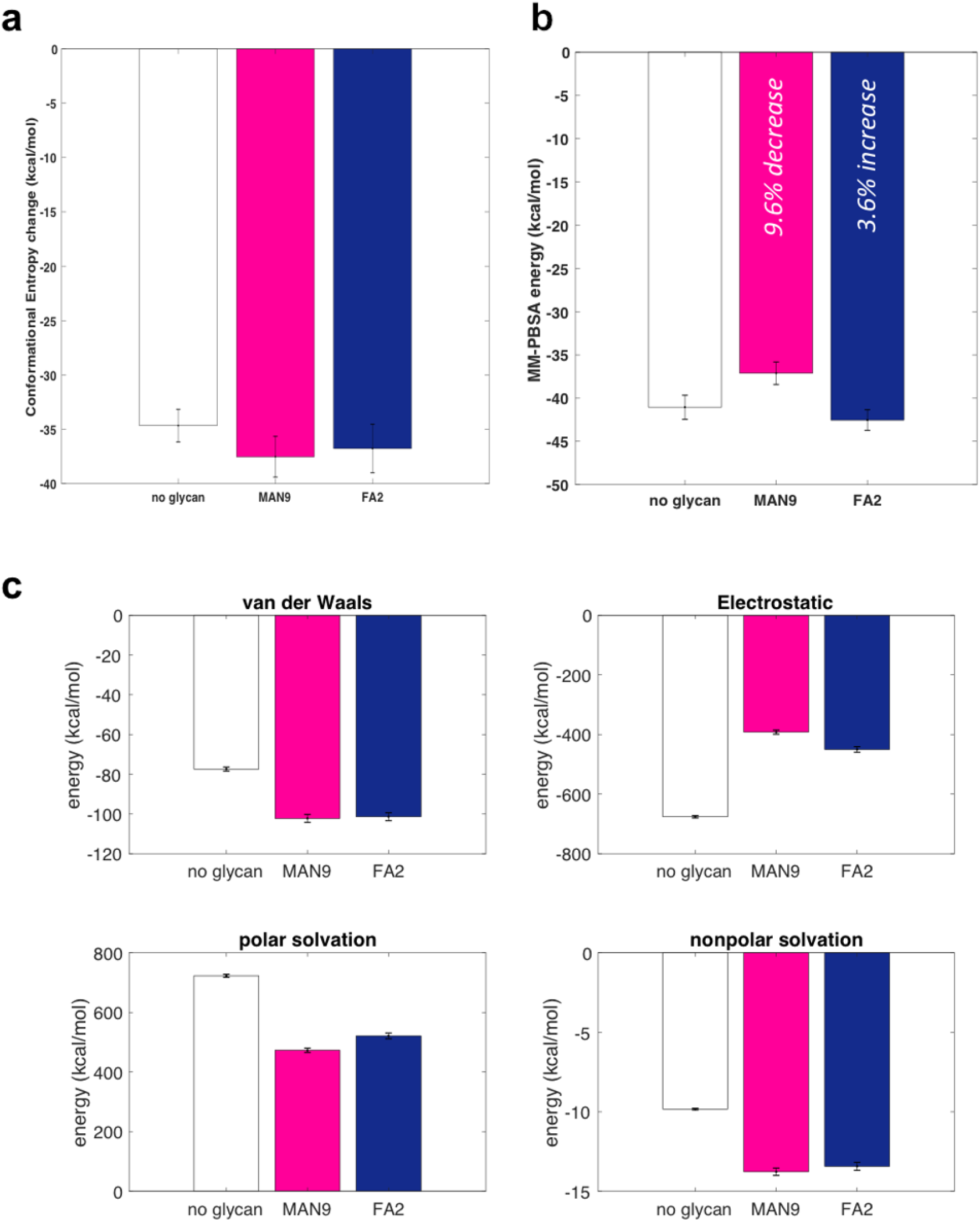
Analysis of RBD-ACE2 binding energy. **(a)** Change in conformational entropy using a quasiharmonic approximation. **(b)** Same MM-PBSA analysis as in main Figure 5a, except that glycan Asn90 is removed from the simulations with MAN9 (magenta), or FA2 glycans (blue) in ACE2. **(c)** Decomposition of contributions to the MM-PBSA energy as shown in main Figure 5a into van der Waals, electrostatic, polar solvation, and non-polar solvation energy components.

**TABLE S1:**
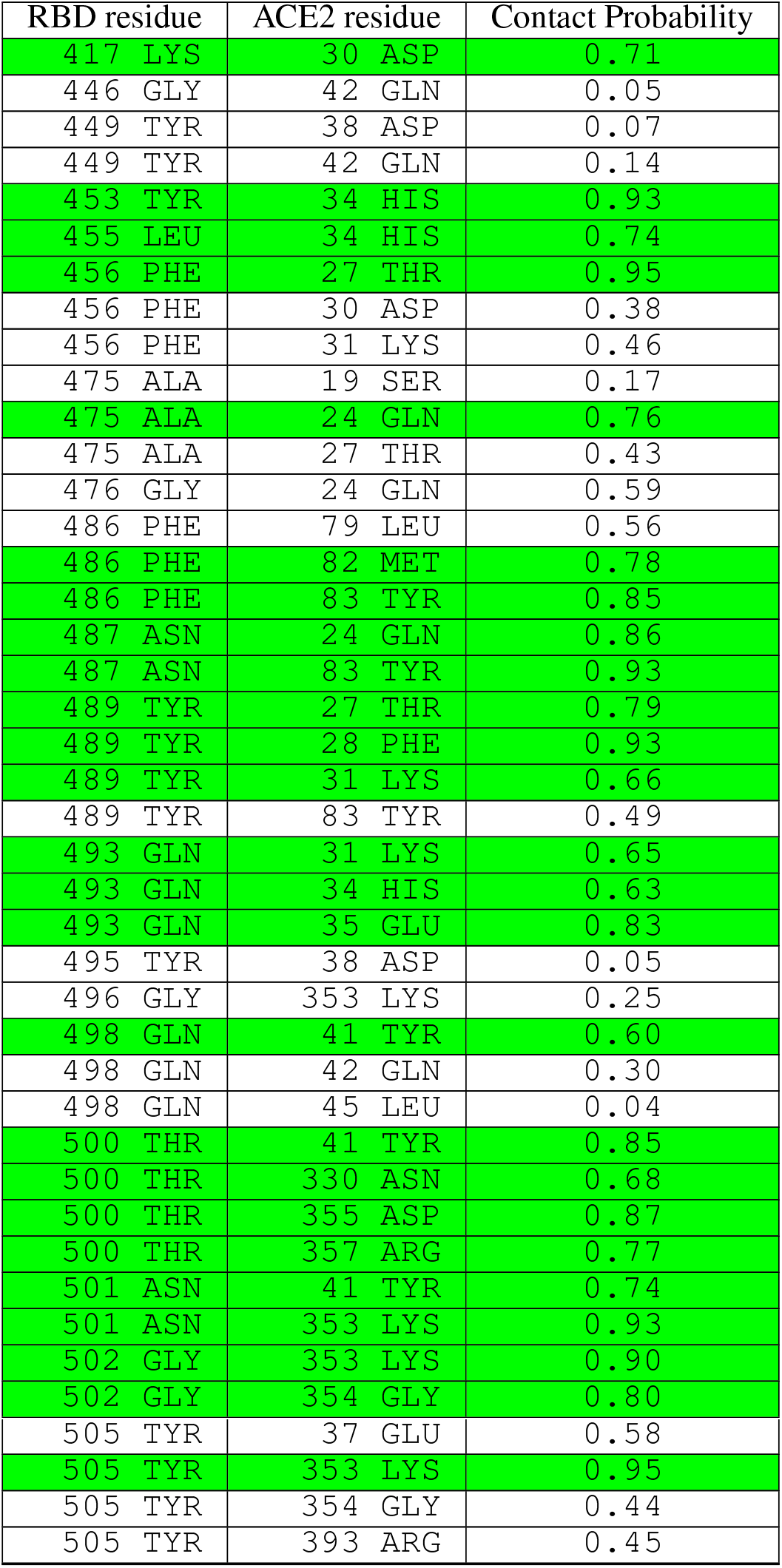
Quantitative evaluation of contacts previously implicated by experimental structures. RBD-ACE2 contacts found in PDB IDs: 6M0J [30], 6M17 [31], and 6VW1 [32] are listed. For each pair, the contact probably was calculated using our allatom simulations. All persistent contacts as captured by the simulations (i.e. total of 25, highlighted in green) represent a subset of the experimentally-reported interactions. Persistent contacts are defined as those that form with at least 60% probability.

